# Detection of a novel mutation G511T in the 530 loop in 16S rRNA in multi drugs resistant Mycobacterium tuberculosis isolated from Sudanese patients

**DOI:** 10.1101/497628

**Authors:** Yousif Mohammed Alfatih, Abeer Babiker Idris, Hadeel Gassim Hassan, Eman O M. Nour, Nihad M A. Elhaj, Salah-Eldin El-Zaki, Muataz M. Eldirdery, Rahma H. Ali, Nuha Y. Ibrahim, Asrar M Elegail, Mohamed Ahmed Salih

## Abstract

**Background:** Tuberculosis (TB) is a bacterial disease considered as a global public health emergency by the World Health Organization (WHO) since 1993. In Sudan, MDR-TB represents a growing threat and one of the most important challenges that faced national tuberculosis program to establish a comprehensive multidrug-resistant tuberculosis management system.

**Objective:** To characterize the diversity and frequency of mutations in Sudanese MDR-TB strains isolated from Wad Madani, Al-Gadarif and Khartoum using 16S rRNA and phylogeny approach.

**Material and Methods:** A total of 60 MDR-TB isolates from Wad-Madani, Al-Gadarif and Khartoum were tested with molecular LPA (Genotype MTBDR plus) and GeneXpert MTB/RIF assay and Spoligotyping to confirm their resistance to RIF and INH. Sequencing and phylogenetic analysis was carried out using in silico tools.

**Result:** This study revealed the circulation of different Sudanese MDR-TB strains isolated from Wad Madani and Al-Gadarif belonging to two distinct common ancestors. Two isolates from Wad Madani (isolate3 and isolate11) found in one main group which characterized by a novel mutation G511T in the 530 loop.

**Conclusion:** The recurrence of C217A mutation in Wad Madani (isolate11) indicates the spread of this mutation in Sudanese MDR-TB strains and the diversity of this inheritance leading to generate new G511T novel mutation. So, understanding the molecular characterization of resistance mechanisms in MD-TB can facilitate the early detection of resistance, the choice of appropriate treatment and ultimately the management of MD-TB transmission. Bioinformatics approaches provide helpful tools for analyzing molecular mechanisms of resistance in pathogens.

## 1. introduction

Tuberculosis (TB) is a bacterial disease considered as a global public health emergency by the World Health Organization (WHO) since 1993 (1–3). It is the ninth leading cause of death worldwide, with a mortality ranging from 1.6 to 2.2 million per year (4, 5), and the leading cause from a single infectious agent, ranking above HIV/AIDS (5–8). However, most deaths from TB could be prevented with early diagnosis and appropriate treatment. According to the global tuberculosis report 2017 which has been published by WHO, In 2016, there were 600 000 new cases with resistance to rifampicin the most effective first-line drug, of which 490 000 had multidrug-resistant TB (MDR-TB) and the treatment remains low globally (5). In Sudan as in other developing countries, MDR-TB represents a growing threat and one of the most important challenges that faced national tuberculosis program to establish a comprehensive multidrug-resistant tuberculosis management system (9, 10).

Multidrug-resistant TB (MDR-TB) is defined as resistance to first-line anti-TB drugs such as Rifampicin (RIF), which is the most effective drug against TB and considered as a broad spectrum antimicrobial (11–14). and isoniazid (INH) drug which has the most powerful bactericidal activity against TB and has good tolerance and low price (14). Multi Drug-resistant Mycobacterium tuberculosis has been reported in the early days of the introduction of drug therapy (15). Grasp the molecular basis of resistance can lead to the development of novel rapid methods for the diagnosis of MDR-TB (16). In addition to highly conserved primer binding sites, 16s ribosomal RNA gene (16S rRNA) sequences contain hyper variable regions that can provide species-specific signature sequences useful for identification of uncharacterized MDR-TB (17, 18). as a result, 16S rRNA gene sequencing has become prevalent in medical microbiology as a rapid and cheap alternative to phenotypic methods of bacterial identification (19, 20). Although it was originally used to identify bacteria, 16S sequencing was subsequently found to be capable of reclassifying bacteria into completely new species or even genera (21, 22). It has also been used to characterize new species that difficult been successfully cultivated in vitro (23, 24).

As long as MDR-TB is not verified, use of inadequate management and hence ineffective antibiotics may lead to further spread of uncharacterized resistant bacteria and amplification of resistance(25, 26). Smear microscopy has long been known as the primary method for screening of TB, with a case detection rate of not more than 68% (27). Marked instance of TB misdiagnosis was evident in 2010 where 2 million of the 5.8 million (34.4%) globally notified cases were found to be smear negative (28). However, the time for bacteriological culture-based diagnosis of TB may require several weeks to months so the rabid and sensitive method will be required for good and safety management (29–31). Therefore, the aims of the current study were to analyze the molecular characterization of Sudanese MDR-TB strains isolated from Al-khartoum, Al-Gadarif and Wad-Madani States using 16S rRNA. Also, to determine the frequency of mutations in 16S rRNA among them. To our knowledge no previous study has reported the sequencing of 16S rRNA in order to study the diversity and frequency of *rrs* mutations among Sudanese strains isolated from Al-Gadarif and Wad-Madani States.

## 2. Material and methods

### 2.1 Ethical consideration

This study was approved by the National center for research ethical committee and the Ethics Committee of the Sudan international University -Khartoum, Sudan.

### 2.2 Study design and study setting

A cross-sectional descriptive study was conducted at the National Public Health Laboratory, Khartoum, Sudan. A total of 60 MDR-TB isolates from Wad-Madani, Al-Gadarif and Khartoum (Figure 1), were tested with molecular LPA (Genotype MTBDRplus) and GeneXpertMTB/RIFassay and Spoligotyping (32, 33) to confirm their resistance RIF and INH. All sample show positive MDR-TB.

**Figure 1.**
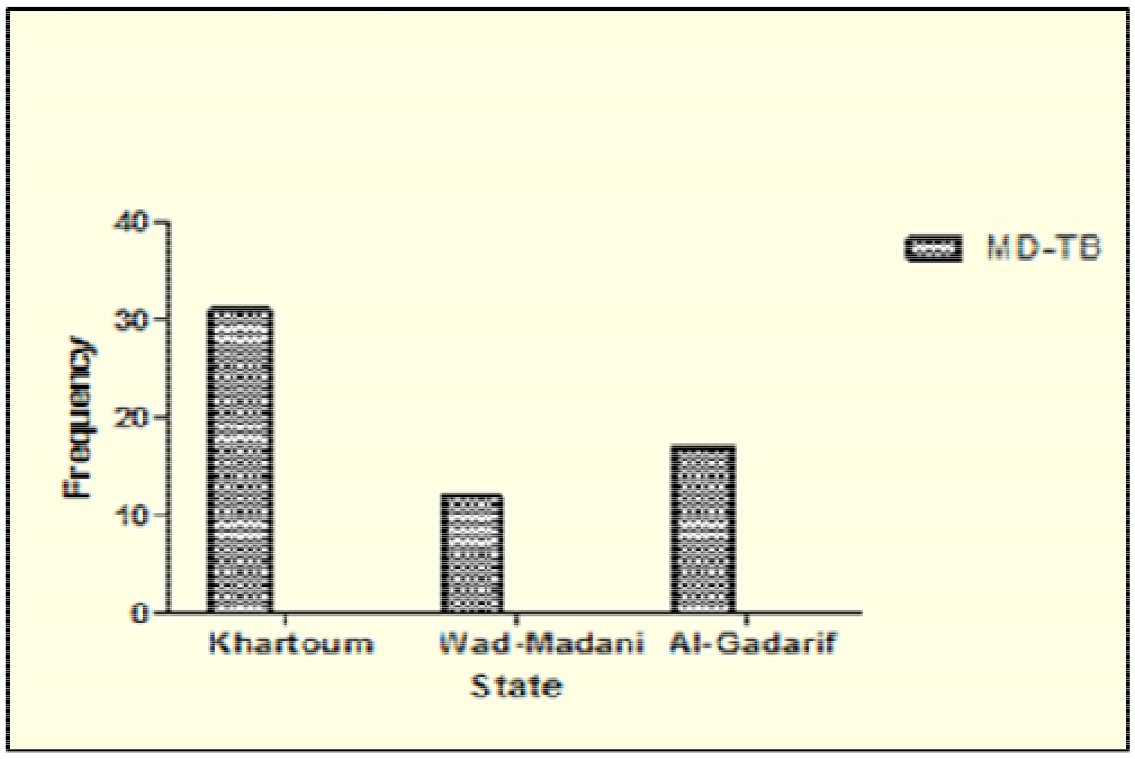
Frequency of our isolates in the states

### 2.3 Molecular analysis

#### 2.3.1 GenoLyse Extraction Method

DNA extraction was performed as recommended by (Hain Lifescience) manufacturer. The first step was to heat the isolate in a water bath for 30 minutes at 10°C and then suspended in100ul of kit lysis buffer followed by incubation at 9°C for 5 min the last step addition of 100ul neutralization buffer. The mixture was spun at full speed in a tabletop centrifuge with an aerosol tight rotor. The extracted DNA were then stored at -2°C (34).

#### 2.3.2 PCR for 16s rRNA gene

Twenty genomic DNA were used as templates for PCR amplification of complete *rrs* gene (16SrRNA). The two primers used were forward primer, 27F(5-AGAGTTTGATCCTGGCTCAG-3) and reverse primer, 1495R(5-CTACGGCTACCTTGTTACGA-3). The 25ul reaction mixture contained 5ul DNA, 2.5ul reaction buffer (10x) with 1.5ul MgCl_2_, 0.5ul Taq DNA polymerase (5 U/ul) 0.625ul dNTPs, 1ul of 10 pmol of each primer, and 1x of gel loading buffer, followed by completing the volume to 12.875ul by DW. PCR amplifying procedure was as follows: Initial Melting (5 min at 94°C), 30 cycles as follow: Melting (1 min at 94°C), Annealing (1 min at 58°C) and Extension (2 min at 72 °C) then final Extension (10 min at 72°C), which was performed on a Bio-Rad (DNA engine/Dyad Peltier) automatic thermal cycler. The products of amplification were checked through running on 0.6% agarose gel electrophoresis (35). The PCR product for *rrs*gene was (1500pb). (Figure 2a)

**Figure 2.**
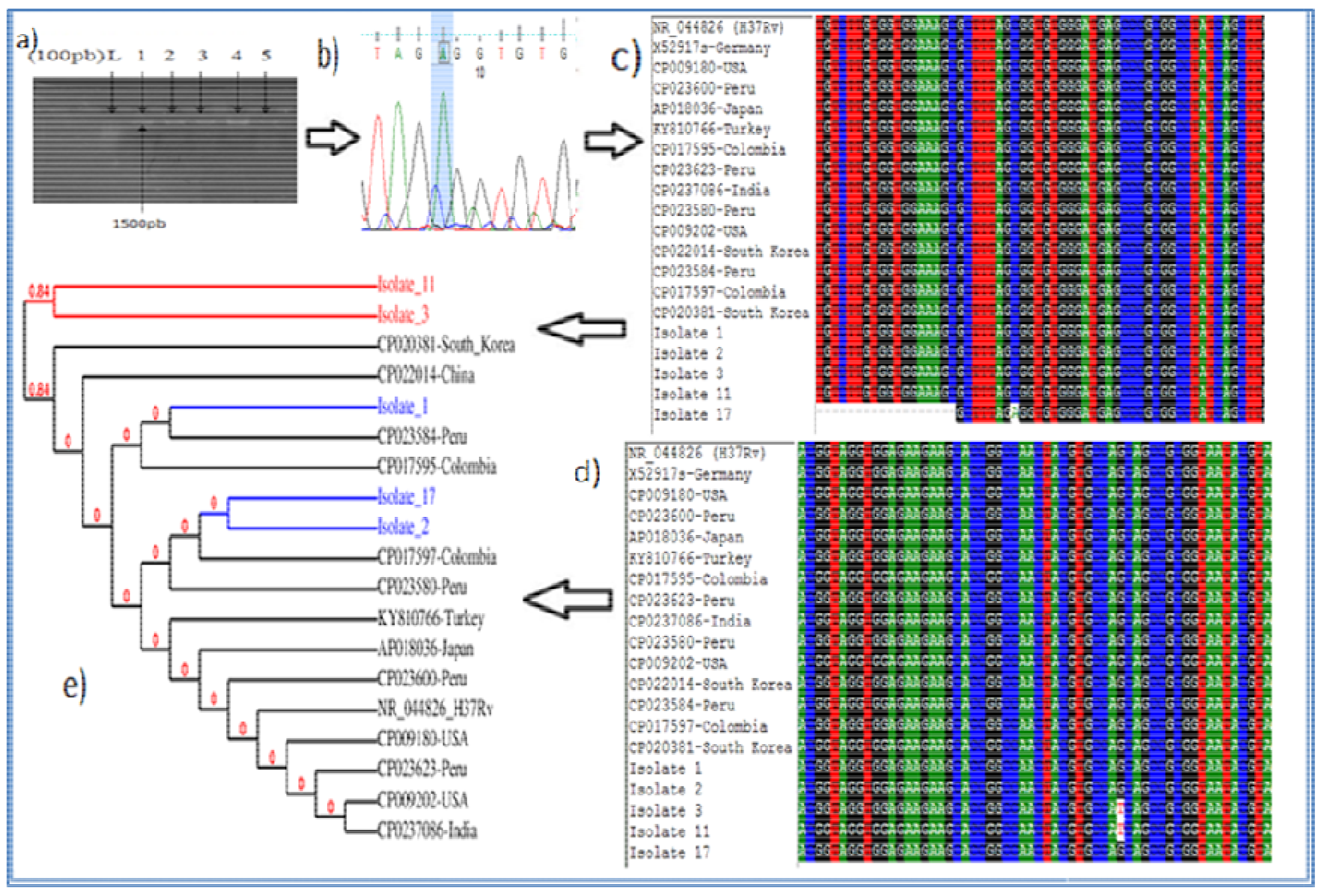
(2a) PCR amplification results of *rrs* gene products on 2% agarose gel electrophoresis. Lane 1 DNA ladder: MW100-1500bp fragments. (2b) Sequencing result of the chromatogram using Finch TV software illustrates C217A mutation. (2c) BioEdit multiple sequences alignment determining C217A mutation. (2d) BioEdit multiple sequences alignment determining a novel mutation at position 511. (2e) Phylogenetic tree using Phylogeny.fr software. The 5 nucleotide sequences of the rrs genes containing novel mutations were deposited in the GenBank database (National Center for Biotechnology Information; https://www.ncbi.nlm.nih.gov/) under the following accession numbers: MG995016, MG995017, MG995018, MG995019, MG995020.

### 2.4 Sequencing of 16S rRNA

PCR products of 15 isolates were packaged according to the International Air Transport Association guidelines and shipped with authorized permission to Macrogen Company (Seoul, South Korea) for BGI Sanger Sequencing. Sequencing for both forward and reverse nucleotide sequencing was carried out.

### 2.5 Bioinformatics analysis

The chromatogram files of sequencing results were viewed and checked for quality by Finch TV program version 1.4.0(36). The Basic Local Alignment Search Tool (BLAST; https://blast.ncbi.nlm.nih.gov/ Blast.cgi)(37) was used to search highly similar sequences available in GenBank database which retrieved from NCBI(38). Then BioEdit software (39) was used to align the similar sequences and the phylogenetic analysis was performed using the online server Phylogeny.fr (http://www.phylogeny.fr/) (40).

### 2.6 statistical Analysis

The results were analyzed statistically using Graph Pad Prism (version 5.0) which is a commercial scientific 2D graphing and statistics software(41).

## 3. Results

### 3.1 Demographic characteristic of patients

Among 60 patients who infected with RIF and INH resistant *M. tuberculosis* isolates, were selected in this study, 32(53.3%) of them were males while females represented 28(46.7%). The highest percentage of MDR-TB was found in the age group ranged between 31-59 years 38(63.3%) while the lowest in the age group ≥60years 3(5%), as illustrated in Table 1.

**Table 1.**
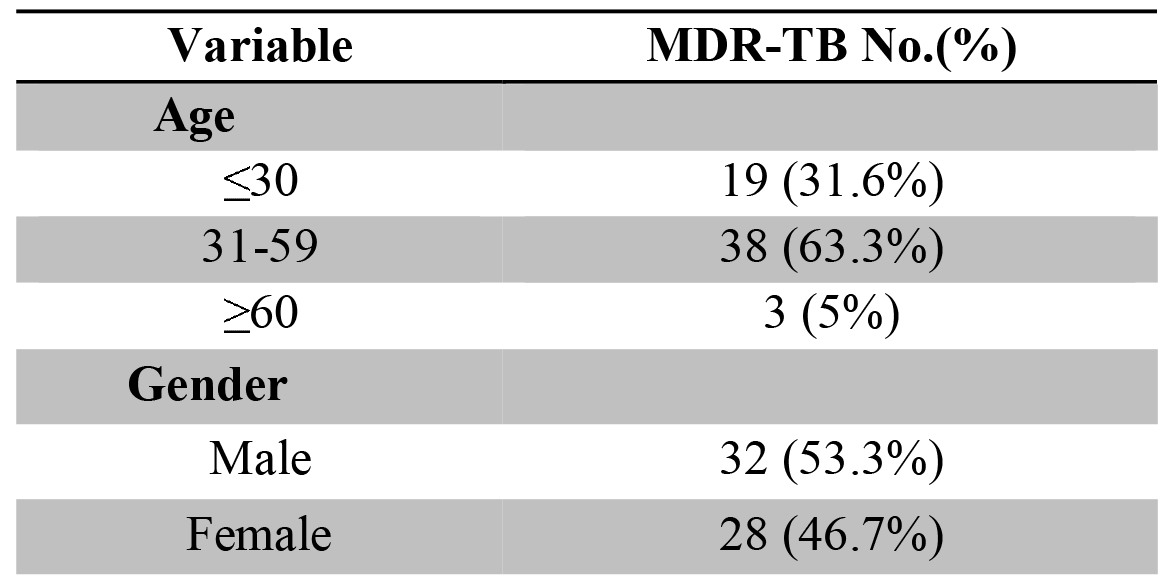
Demographic characteristics of MDR-TB infected patients

### 3.2 Bioinformatics analysis result

From 15 chromatograms, only five showed high quality by FinchTV. BLAST analysis of their sequences showed 99% identity to 16s rRNA gene nucleotide sequences of different MDR-TB strains isolated from different countries, that obtained from GenBank databases, accession numbers of 16s rRNA gene nucleotide sequences of MDR-TB retrieved strains and their area of collections are listed in Table 2.

**Table 2.** Accession numbers of *rrs* gene nucleotide sequences of MDR-TB retrieved strains and their area of collections

| **Gene Bank nucleotides**<br>**accession No.** | **Country** |
| --- | --- |
| **CP022014** | China |
| **CP020381** | south-Korea |
| **CP0235801** | Peru |
| **CP017597** | Colombia |
| **KY810766** | Turkey |
| **AP018036** | Japan |
| **CP009180** | USA |
| **CP009202** | USA |
| **CP0237086** | India |
| **X52917** | Germany |

The multiple sequence alignment of the 5 Sudanese MDR-TB *rrs* genes nucleotide sequences with the similar nucleotide sequences that obtained from BLAST was carried out, to find the homology and evolutionary relation between these sequences, by BioEdit software; after checking the identities and differences we found that two isolates from Wad Madani (isolate3 and isolate11) had a novel mutation, G→T transversion at nucleoside position 511, in the 530 loop. Al-gadarif’s isolate (isolate17) showed C →A transversion at nucleoside position 217. Another novel mutation, G→A transition at nucleoside position148, was found in two isolates (isolate2 and isolate3) from Wad Madani. However, isolate3 harbored another third novel mutation, G →T transversion at nucleotide position 736. The phylogenetic tree assimilates and represents the result of relationship between the Sudanese isolates and GenBank series of Mycobacterium tuberculosis stains (Figure 2).

## 4 Discussions

The 16S rRNA is the most common housekeeping genetic marker, so its sequences are used to study bacterial phylogeny and taxonomy (42). In this study, we used the 16S rRNA sequencing in order to characterize the diversity and frequency of mutations in Sudanese strains isolated from Wad Madani and Al-gadarif which revealed the circulation of different strains belonging to two distinct common ancestors. Two isolates from Wad Madani (isolate3 and isolate11) found in one main group which characterized by a novel mutation G511T in the 530 loop. while other isolates from Wad Madani with Al-gadarif’s isolate (isolate17) were clustered in one group and showed close similarity to strains from different part of the world (Figure 2e)

Streptomycin was the first aminoglycoside which discovered in the 1940s. It inhibits initiation of mRNA translation.(43) The rapid emergence of resistant mutants with relatively high frequency among microorganisms has greatly reduced its therapeutic effectiveness. So, Streptomycin should not be used alone to treat any infection. it is used in combination with other anti-tuberculous drugs (INH, rifampin) to treat tuberculosis. (44) In tuberculosis, the high level of resistance against Streptomycin results from missense mutations in the genes *rrs* and *rpsL* which encoding two components of the ribosome, the 16S rRNA and the S12 protein, respectively.(45) when streptomycin-resistant strains with wild-type S12 proteins were found, the resistance has been attributed to mutations in *rrs* affecting one of two highly conserved loops that are adjacent in the 16S rRNA, the regions around nucleotide 904 and the 530 loop.(46) However, about one-third of streptomycin-resistant isolates do not have mutations in the *rrs* or *rpsL* genes.(47)

In this study, we found a novel mutation, G→T transversion at nucleoside position 511, in the 530 loop while alterations at positions 491, 512, 516 were not found.(48–50) Also, the insertion of cytosine residue between nucleotides 512 and 513, which had previously been reported to confer both streptomycin resistance and dependence, was not detected.(6) However, this finding is in agreement with another study done by Katsukawa *et al.* in Japan which found a novel mutation, at that time, at position 513 (A:C, A:T) and also they found no mutations at positions 491, 512, 516, 904, 905 or base insertion between 512 and 513.(47) in contrast, there are another studies conducted in Khartoum state by Solima *et al.(51)*, in Poland by Brzostek et *al.(52)* and in New York City by Cooksey *et* al.(43) revealed no mutations in the 530 loop. Another novel mutation, G→A transition at nucleoside position148, was found in two isolates (isolate2 and isolate3) from Wad Madani. It is interesting to note that isolate3 harbored another third novel mutation, G→T transversion at nucleotide position 736, so it belonged to distinct common ancestor along with isolate11 that had one of these novel mutations G115T. Further studies are needed for in vitro phenotypic resistance with large sample size to confirm the association between these novel mutations and phenotypic Streptomycin-resistance and reliability of these mutations as molecular markers of resistance.

Moreover, Al-gadarif’s isolate showed C217A mutation which is the same novel mutation reported from Streptomycin resistance Sudanese strains in Khartoum state.(51) Regarding the region centered around nucleotide 912, which may form part of a site involved in codon-anti-codon interaction and proofreading(49), there is no mutation was observed which is in accordance with other studies conducted in Barcelona,(53) China,(54) Germany(49) and in Latvia.(55) This accordance could indicate that mutations in the *rrs* 912 region are not common.

## Conclusion

The recurrence of C217A mutation in Wad Madani (isolate11) indicate the spread of this mutation in Sudanese MDR-TB strains and the diversity of this inheritance leading to generate new G511T novel mutation. So, understanding the molecular characterization of resistance mechanisms in MD-TB can facilitate the early detection of resistance, the choice of appropriate treatment and ultimately the management of MD-TB transmission. Bioinformatics approaches provide helpful tools for analyzing molecular mechanisms of resistance in pathogens.

## Acknowledgments

The authors are grateful to Tropical Medicine Research Institute, Total Lab care Laboratories, Molecular Tuberculosis Reference Laboratory, Africa City of Technology, and all members of the National Reference Laboratory of Tuberculosis (NRL-TB), Sudan, Khartoum.

